# Integrative Transcriptomic Analysis Reveals Differential Exon Usage and Virulence Gene Family Dynamics in *Plasmodium falciparum* during Severe Malaria

**DOI:** 10.64898/2026.11.06.765005

**Authors:** Prince Mensah Ansah, Welbeck Odame Dzadey, Alfred Martey, Harun Abdullah, Wisdom Awuku

**Affiliations:** Department of Biochemistry and Biotechnology, Kwame Nkrumah University of Science and Technology, Kumasi, Ghana; Kumasi Centre for Collaborative Research in Tropical Medicine (KCCR), Kumasi, Ghana; University of Missouri, Kansas City, USA; Department of Chemistry and Biochemistry, Auburn University, Auburn, Alabama, USA

## Abstract

Malaria remains a leading cause of child mortality in developing countries, with roughly 610,000 deaths per year. While the disease is caused by five human Plasmodium species, *Plasmodium falciparum* (*Pf*) causes the most severe form. Progression from uncomplicated to severe malaria is characterized by high proliferation of the parasite, severely affecting children under five years old. However, the molecular mechanisms underlying disease progression remain an active area of research. Many multicopy genes have been implicated in *Pf* virulence. Some of these genes are implicated in cytoadherence, host-cell invasion, immune evasion, sequestration of infected erythrocytes, and antigenic variation. We employed an integrative transcriptomic analysis in this study to elucidate the molecular underpinnings of severe malaria and gene expression patterns, virulence gene dynamics, and the potential contribution of alternative splicing.

Results show an overexpression of protein synthesis and ribosomal genes associated with severe malaria. About 86% (n=1865) of all differentially expressed genes (DEGs) were downregulated in severe malaria, and 226 genes exhibited both differential expression and differential exon usage (DEU). *Pf* virulence genes and antigenic genes such as *var* (pfEMP1), *stevor*, *surfin*, *rif* (rifin), *ptm-2TM*, and *msp* were upregulated in severe malaria compared with uncomplicated malaria, except for *rifin*. In severe malaria, there was evidence of co-expression and regulation within multigene families, as well as DEU and alternative splicing in virulence genes and antimalarial resistance-related proteins.

*Pf* does not employ a single mechanism for disease progression but suppresses and deregulates nonessential pathways, processes, and genes, which are upregulated virulence genes, DNA replication, mRNA transcription, and protein synthesis. Multicopy genes are co-expressed and regulated to meet the molecular needs of the parasite. The parasite employs alternative splicing and DEU mechanisms for immune invasion and evasion, host-parasite interaction, as well as antimalarial resistance.

**Author Summary:** Our findings reveal distinct transcriptional patterns between severe and uncomplicated malaria, including differential expression of key parasite virulence-associated gene families. Notably, our analysis highlights differential regulation of the var (PfEMP1), msp (merozoite surface protein), and rifin gene families, suggesting that parasite adhesion, host–parasite interactions, and immune modulation contribute differently to malaria severity. Also, identifying differential exon usage adds another layer of insight into the transcriptomic complexity of P. falciparum and suggests potential regulatory mechanisms that may contribute to parasite adaptation and pathogenicity. We believe these findings advance our understanding of the molecular mechanisms underlying *P. falciparum* pathogenicity through an integration of transcriptional regulation, exon-level variation, and virulence-associated gene family dynamics. By providing a broader perspective on the parasite’s transcriptional responses associated with severe malaria, our findings may inform future investigations into parasite biology, host–pathogen interactions, and the development of strategies for malaria intervention.

## Introduction

Despite multifaceted control strategies, the burden of malaria on public health systems persists. In 2024, an estimated 282 million cases of malaria were recorded globally and 610,000 malaria-related deaths, an increase of about 9 million (3%) from the previous year, with the African region contributing to 95% of cases and mortality of the global burden (Venkatesan, 2025). The causative organism of the disease is the protozoan Plasmodium. Currently, there are five known species, *Plasmodium falciparum, Plasmodium malariae, Plasmodium ovale, Plasmodium vivax*, and *Plasmodium knowlesi*, that infect humans. Of these, *P. falciparum* (Pf) is the most virulent, with its burden higher in children under five years of age and pregnant women. *Pf* pathogenicity is linked to its ability to invade erythrocytes across a broad range of maturation stages, achieve high parasitaemia, and induce the cytoadherence of infected erythrocytes to blood vessels, resulting in sequestration and microvascular obstruction. These processes contribute to serious complications such as acute kidney injury, pulmonary oedema, severe anaemia, haemoglobinuria, circulatory collapse, and multiorgan failure (Chellappan & Bhadauria, 2016; Talwar *et al*., 2019; White, 2022).

Virulence gene families in *Plasmodium falciparum* (Pf) contribute to parasite survival and pathogenicity and are implicated in severe malaria complications. The sequenced Plasmodium genome harbours an extensive, often species-specific array of variant surface antigens (VSAs), which is encoded by the Plasmodium 3D7 reference genome (Cunningham *et al*., 2010). The Plasmodium VSA gene family includes *var*, *rifin*, and *stevor*, which encode erythrocyte surface proteins known as *Pf* erythrocyte membrane protein 1 (PfEMP1), *Plasmodium falciparum*-encoded repetitive interspersed families of polypeptides (RIFIN), and subtelomeric variant open reading frame (STEVOR), respectively. Most clinical studies on the Plasmodium VSA gene family have focused on the var (PfEMP1) family rather than the other families. Malaria infection is achieved by the adherence of malaria-infected erythrocytes to uninfected erythrocytes and endothelial cells, to other healthy red blood cells (resetting), and to leukocytes/serum proteins, resulting in severe clinical complications (Frech & Chen, 2013; Wahlgren *et al*., 2017).

These genes regularly undergo antigenic variation and changes in gene expression, enabling them to evade the host immune system and establish chronic infection (Obi *et al*., 2025; Su *et al*., 1995). Although PfEMP1 has been implicated in severe malaria, the dynamics of this surface adhesion protein and other proteins involved in disease progression are understudied. Recent studies suggest that involvement of non-3d7 VSAs, such as fikk, phist, and surf (Cummings *et al*., 2025), and Peptide VAR P5 (Bhandari *et al*., 2021) are implicated in severe malaria and overall disease progression. This may vary across different epidemiological zones and levels of endemicity. Understanding the patterns could be useful for informing the design of vaccines targeting severe malaria disease.

Due to limited data on the association of Pf field isolates with different malaria clinical manifestations, that is, asymptomatic, uncomplicated, and severe malaria, our understanding of the biological processes underlying *Pf* virulence is limited. Thus, the relationship between parasite genetic profiles and malaria-related pathologies needs to be established to better understand the pathogenesis of the disease.

We used an integrative transcriptome approach in this study to identify *Pf* genes differentially expressed between uncomplicated and severe malaria. We further identified genes that undergo both differential expression and exon usage during disease progression. Lastly, we assessed the dynamics of the plasmodial virulence gene families.

## Results

### Differential Gene Expression

Differential gene expression analysis identified 1,849 (34.3%) of the 5,396 *Plasmodium* genes assessed as differentially expressed. Of these differentially expressed genes (DEGs), 258 (13.95%) were upregulated, whereas 1,591 (86.05%) were downregulated, indicating high transcriptional downregulation in severe malaria (**<u>Figure 1a, b</u>**).

**Figure 1.**
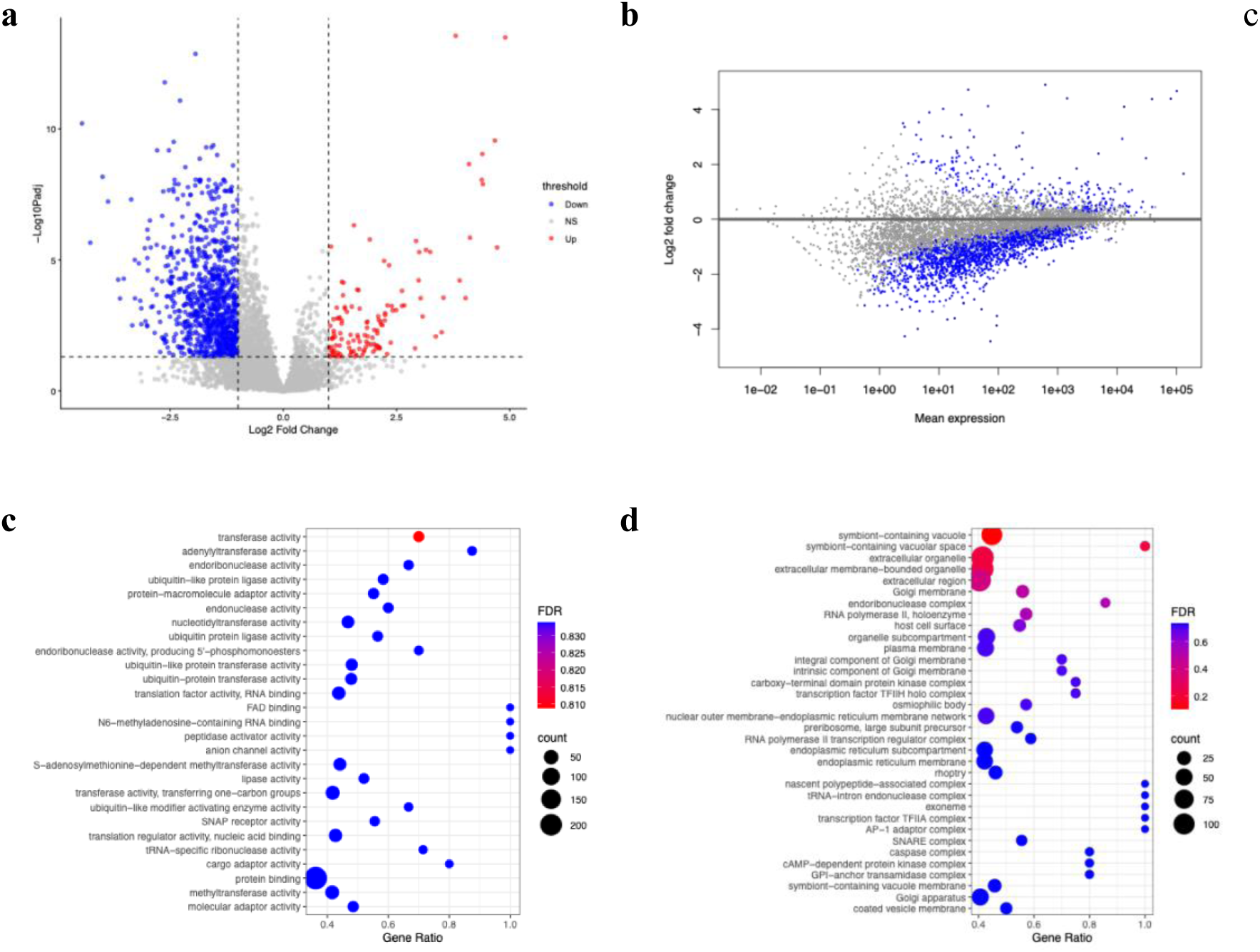
Differential gene expression analysis. (a) Volcano plot showing differentially expressed genes. (b) MA plot showing differential gene expression. (c–d) Gene Ontology enrichment analysis of differentially expressed genes for molecular function and cellular component terms.

Gene Ontology (GO) enrichment analysis and enrichment profiles suggest that the identified genes were mainly involved in enzymatic modification processes, RNA metabolism, protein regulation, and intracellular signaling mechanisms (**<u>Figure 1</u>**). Among the molecular functions, transferase activity and adenylyltransferase activity had the highest gene ratio and enrichment significance. FAD binding, N6-methyladenosine-containing RNA binding, peptidase activator activity, and anion channel activity, although they showed the highest gene ratio, had lower counts. Protein binding activity in the enriched molecular functions showed the highest counts, although with the lowest gene ratio. Additional enriched molecular functions included endonuclease activity, endoribonuclease activity, DNA binding, cargo adaptor activity, and molecular adaptor activity, with cargo adaptor activity showing a high gene ratio. Enrichment of ubiquitin-protein transferase activity, ubiquitin-protein ligase activity, ubiquitin-like protein ligase activity, ubiquitin-like protein transferase activity, and ubiquitin-like modifier activating enzyme activity suggested the involvement of protein ubiquitination and related post-translational modification processes.

GO cellular component enrichment analysis revealed significant enrichments in genes associated with intracellular trafficking, membrane-bound organelles, and transcriptional regulatory complexes (**Figure 1**). In severe malaria, the symbiont-containing vacuole and symbiont-containing vacuolar space showed significantly higher enrichment levels in *Pf* genes than in uncomplicated malaria.

Extracellular organelle, extracellular membrane-bounded organelle, and extracellular region were also significantly enriched, despite involving relatively few genes. Genes associated with transcription machinery, such as RNA polymerase II holoenzyme, RNA polymerase II transcription regulator complex, transcription factor TFIIH holo complex, and transcription factor TFIIA complex, were enriched in severe malaria relative to uncomplicated malaria. Additionally, genes linked to intracellular transport, such as Golgi membrane, Golgi apparatus, Integral component of Golgi membrane, Intrinsic component of Golgi membrane, and coated vesicle membrane, were enriched. Other genes linked with cellular components, such as nascent polypeptide-associated complex, tRNA-intron endonuclease complex, exoneme, and AP-1 adaptor complex, had the highest gene ratio, suggesting high specific enrichment of these genes. Genes linked to the endoplasmic network, such as nuclear outer membrane-endoplasmic reticulum membrane network, endoplasmic reticulum membrane, and endoplasmic reticulum subcompartment, were enriched in the GO cellular component enrichment analysis.

### Differential Exon Usage (DEU)

Differential exon usage was detected in 834 genes, indicating changes in the relative usage of individual exons within these genes (**Figure 2**). Of these, 226 genes also showed significant differential gene expression, suggesting that both transcript abundance and exon usage were altered in this subset of genes (**Figure 3**). Out of the 225 genes undergoing both alternative splicing and differential expression in tandem, 124 were upregulated, while 102 were downregulated. *PfEMP1* and genes encoding ribosomal proteins show active exon usage in severe conditions. The top 5 genes with high DEU include chloroquine resistance transporter (PDF_0709000), nuclear polyadenylated RNA-binding protein NAB2 (PDF_0623100), metacaspase-2 (PF3D7_1438400), conserved plasmodium protein (PF3D7_0713700), and GTP-binding protein (PFD7_1358900) (**Figure 2**). These genes are implicated in sporogony and gametogenesis of Plasmodium parasites. They are present in the parasite but absent in the host, making them potential targets for anti-malaria drug development (Kumari *et al*., 2022).

**Figure 2.**
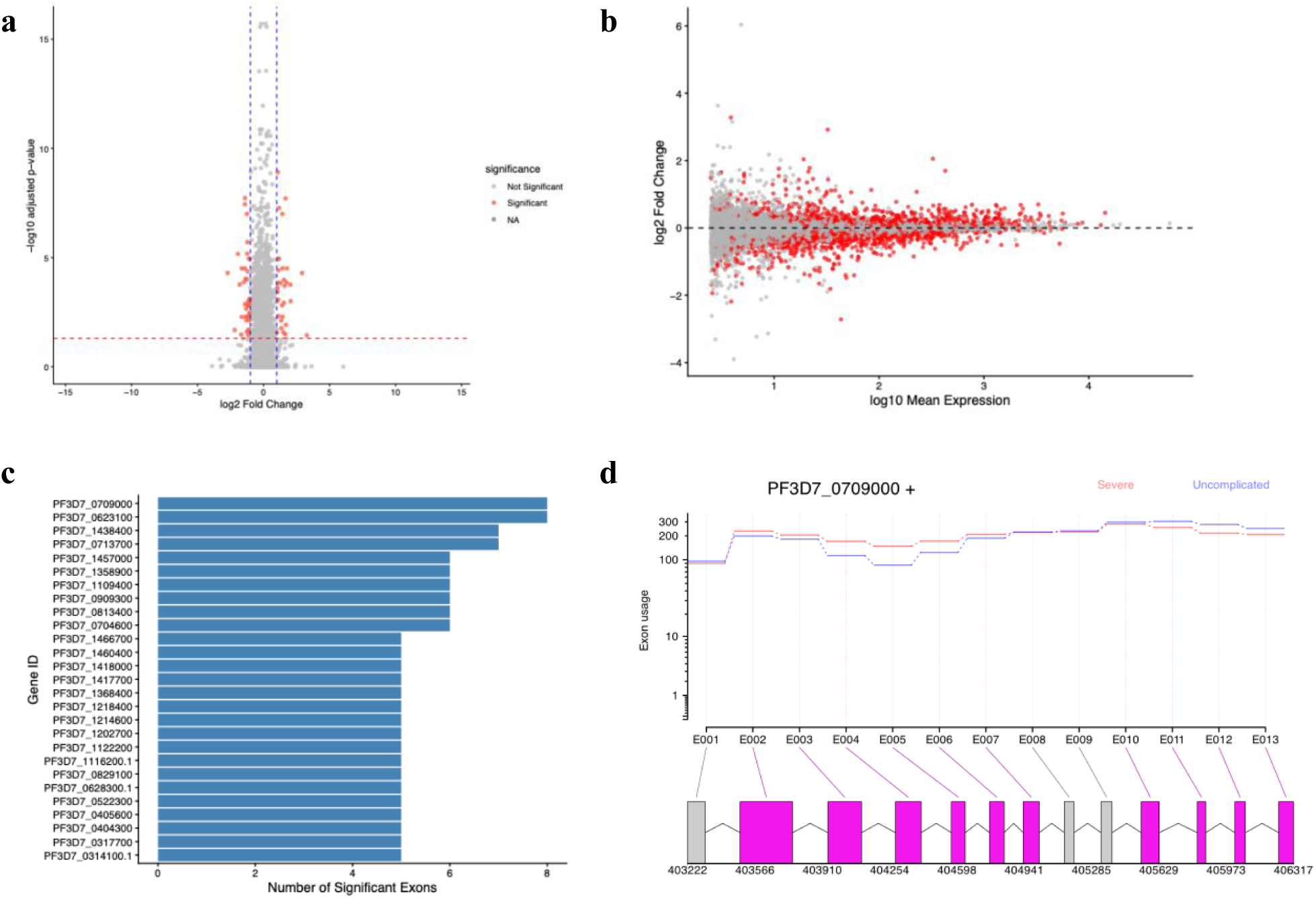
Differential exon usage. (a) Volcano plot showing differentially used exons. (b) MA plot showing differential exon usage. (c) Bar plot of the top genes exhibiting differential exon usage. (d) Transcript plot of Differential exon usage of chloroquine transporter protein (PF3D3_0709000)

**Figure 3.**
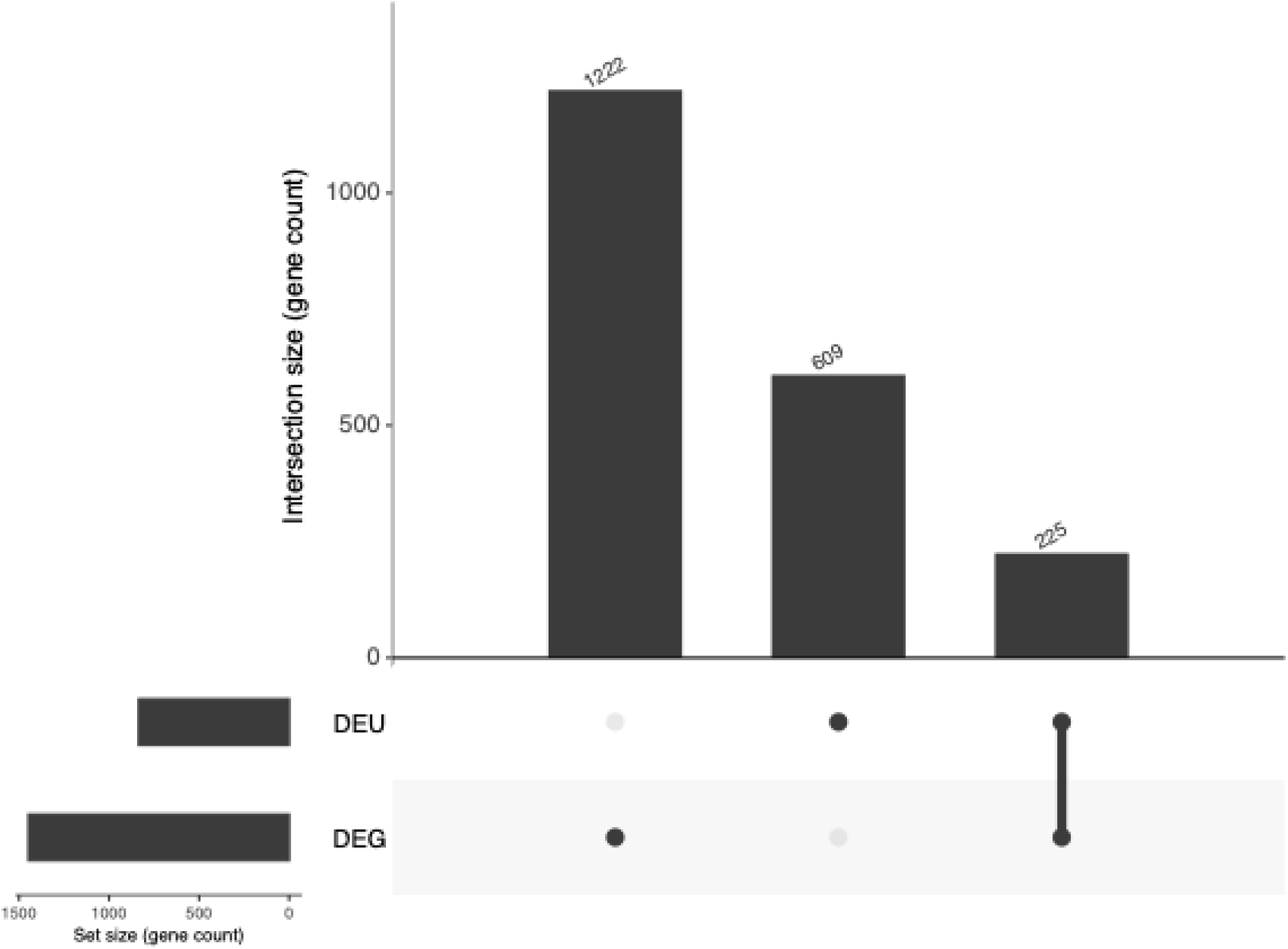
Differential gene expression and exon usage. UpSet plot showing the overlap between genes exhibiting differential expression and differential exon usage. Genes in the overlap represent candidates with coordinated changes in transcript abundance and exon usage.

**Figure 4.**
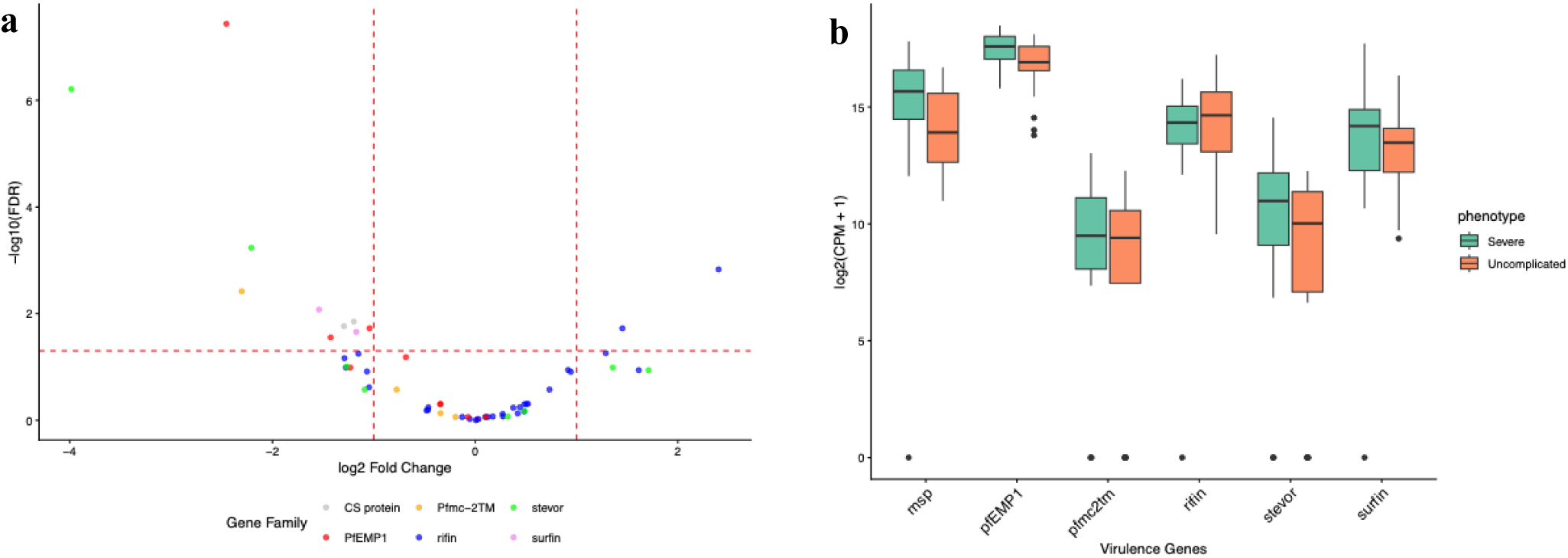
Plasmodium *falciparum* virulence gene family expression dynamics. Expression patterns of pfEMP1, *rifin, Pfmc-2TM, msp, stevor, and surfin* gene families in severe and uncomplicated malaria.

### Virulence Gene Dynamics

Virulence-associated gene expression patterns were assessed by compiling genes previously implicated in malaria severity, including those involved in cytoadherence and parasite–host cell interactions. Overall, virulence-associated genes showed higher expression in severe malaria compared with uncomplicated malaria. Higher expression levels were observed among *pfEMP1* and *msp* gene families. In contrast, *rifin* genes exhibited an inverse expression pattern, with relatively higher expression in uncomplicated malaria than in severe malaria (**Figure 3**). The expression patterns of virulence genes differ markedly between severe and uncomplicated malaria (**<u>Figures S2</u> and <u>S3</u>**). There is an observation of co-expression and regulation among the various isoforms of members of the gene family.

Overall, genes within the same family exhibited strong positive correlations in their expression patterns, indicating coordinated expression among family members. However, a few gene families showed inverse expression patterns, with increased expression of some family members associated with reduced expression of others (**<u>Figure S4</u>**).

## Discussion

Malaria severity is driven by molecular signatures in the infectious organism’s genome. Severe malaria is characterized by massive parasite proliferation leading to high parasitemia. The gene expression analysis highlighted dysregulated genes that could serve as molecular signatures during disease progression. A large proportion of highly differentially expressed genes have functions in protein synthesis and ribosomal assembly. Protein synthesis is a function of the ribosome and its accessory proteins. Thus, the upregulation of these genes is quite expected. A high rate of proliferation must be commensurate with the rate of protein synthesis. Malaria severity is also affected by the host immune response and parasite-host interactions (PHI). This interaction inadvertently affects the parasite’s overall survival in the host. This observation is highly corroborated by the enrichment of pathways leading to protein synthesis. Gene ontology analysis reveals the enrichment of protein synthesis molecular terms and cellular components. To upregulate protein synthesis, the transcriptional machinery needs tuning to feed into protein translation. Hitherto, this is seen in the dysregulation of certain transcriptional accessory proteins and transcriptional factor modification.

The role of *Pf* multigene families in antigen-mediated immune evasion in infected red blood cells has been reported. Niang *et al*. showed, using a combination of immunofluorescence assay (IFA), flow cytometry, and agglutination tests, that the *stevor* multigene family modifies Plasmodium-infected red blood cells (iRBCs) (Niang *et al*., 2009; Omelianczyk *et al*., 2020). The expression of *stevor* on iRBC surfaces facilitates changes in antigenic properties. The high expression of the *stevor* multigene family in this study supports it as a key cluster of proteins needed for host-parasite interactions (Niang *et al*., 2009). The relative expression of *stevor* multigene proteins is high in severe malaria compared to uncomplicated malaria. Interestingly, some studies found that the levels of *stevor* expression also differ between patient-isolated *Pf* and lab-cultured isolates (Blythe *et al*., 2008). This is quite expected, given the role of *stevor* in the parasite’s pathobiology. Specific subtypes of *rifin* offer both clinical immunity and immune evasion mechanisms, securing their role in severe malaria pathogenesis (Kassegne *et al*., 2025). In this study, the rate of expression of *rifin* is counterintuitive. *Rifin* has been associated with reduced risk of severe malaria, facilitation of immune suppression, and mimicry. *Rifin* is associated with reduced risk of severe malaria, supporting the reversal in expression pattern among the multi-gene families. In Figure 3, the rate of expression of *rifin* in uncomplicated malaria is relatively higher than in severe malaria. *Rifin* has been extensively characterized through immunoproteomic assays. The presence of the LILRBI motif presents a vaccine target for next-generation anti-malaria agents to combat severe malaria and reduce the global burden of malaria (Kassegne *et al*., 2025).

PfEMP1 genes are pivotal molecular markers of cytoadherence, which are required for parasite invasion and proliferation; thus, *Pf* makes several copies of these genes. More importantly, the results from this study show that they also undergo extensive alternative splicing.

Overall, these virulence-associated gene families exhibited distinct expression patterns between the two clinical groups, with *msp* showing relatively higher expression in severe malaria, while *rifin* displayed an inverse pattern with relatively higher expression in uncomplicated malaria. Virulence gene families are highly expressed in severe malaria compared to uncomplicated malaria, except *rifin*. This makes sense because it seems *Pf* suppresses nonessential processes and upregulates virulence families. The findings from this study also highlight the gene family dynamics. Essentially, the members of each multigene family are not expressed or spliced in the same pattern. Importantly, there is evidence of co-expression and co-regulation within the multi-gene family. This shows that a subset of the multi-gene family may be involved in different molecular tasks. Evidence of this is seen in the rifin multigene family, where a subset of members provides inhibitory and relieving roles in severe malaria (Andersson *et al*., 2020; Bachmann *et al*., 2015; Kassegne *et al*., 2025; Niang *et al*., 2009). It is interesting to point out that generally, most of the genes are co-regulated. The positive correlation shows agreement in molecular role within the multigene copy. Conversely, genes that are negatively co-regulated may have opposing molecular roles regarding severity of malaria.

Alternative splicing is another method the organism uses to efficiently enforce proliferation in the host organism. Additionally, alternative splicing of key proteins helps the parasite evade its host immune system. It seems the organism may employ this same method for developing an arsenal against antimalarials (See <u>Figure 2</u>). Chloroquine transporter protein and metacaspase-2 are undergoing massive alternative splicing (differential exon usage). This is an interesting finding pointing to one of the molecular underpinnings of antimalarial drug resistance. This study reports direct evidence of DEU and alternative splicing in key virulence genes and protein synthesis machinery driving the severity and disease progression. The use of alternative splicing by organisms as a means of survival is not uncommon. We report that not only does the Plasmodium parasite employ overexpression of antigenic variation in its parasite pathobiology, but it could also potentially use alternative splicing as a molecular mechanism to evade the host immune system and express various antigens on the iRBCs. Chloroquine transporter protein and metacaspase-2 are under significant DEU. This finding, among others, points to another molecular mechanism of anti-malaria resistance development. The targeted protein becomes resistant to the drug target through active protein structural metamorphosis driven by alternative splicing forces. The unending combinatorial protein antigenic synthesis through different exon combinations makes Plasmodium indefinitely insusceptible to antimalarial drugs. It also enhances the active adaptation of the parasite to antimalarial agents. Metacapsases are implicated in sporozoite proliferation and chloroquine-resistant transporters (Kumari *et al*., 2022). *Pf* has multiple copies of these hypervariable genes. More importantly, they undergo extensive alternative splicing and are also overexpressed, giving the organism a robust force to drive disease progression and severity.

## Conclusion

This study corroborates the molecular basis of malaria severity and disease progression through integrative transcriptomic analysis. *Pf* employs the downregulation of competing pathways and molecular processes while upregulating genes involved in protein synthesis, parasite-host interaction, cytoadherence, immune evasion, and modulation. Here we report, for the first time, evidence of differential exon usage and alternative splicing in severe malaria. Alternative splicing could potentially be a means of evading the host immune system and a molecular mechanism for developing resistance to anti-malaria agents. We report the dynamics of virulence gene families serving as hypervariable antigenic variation targets. The results rehash and highlight the role of these essential genes as molecular signatures and targets for potential vaccine and anti-malaria drug development.

## Study Limitation

The study used existing data, which means we did not have full control of the data collection and conditions for data collection. We propose that independent samples should be collected to validate the DEU signal. Also, most of the genes in the *Pf* genome are annotated as putative. This lack of experimental data to corroborate the computationally predicted proteins makes it difficult to use them as targets.

## Materials and Methods

### Genomic Data Acquisition, Data QC, and Read Mapping

To assess the differential expression and exon usage dynamics, we downloaded transcriptomic data from the NCBI database (project ID PRJEB21707) using the NCBI SRA Toolkit. The data consisted of *Pf* isolates from severe (n=23) and uncomplicated (n=21) malaria cases (Table S1). The Plasmodium *falciparum* genome (GCF_000002765.6) and annotation file were downloaded from NCBI and assessed for completeness and quality.

FastQC 0.12.1 (Andrews, 2017) and MultiQC (Ewels *et al*., 2016) were used to assess read quality. To ensure uniformity in data quality across samples, fastp (Chen *et al*., 2018) was used for trimming with default settings and *w 4: --detect_adapter_for_pe --overrepresentation analysis --dedup --dup_calc_accuracy 3 --overrepresentation_sampling 10000 --trim_front2 12 --trim_front1 12*. Reads were aligned to the Plasmodium *falciparum* reference genome using STAR with the *--quantMode* option enabled to generate transcript-level read counts for downstream gene expression analysis (Dobin *et al*., 2013).

### Differential Gene Expression Analysis

Differential gene expression analysis was performed using DESeq2. We filtered out genes with low read counts (<20) before downstream analysis. Differentially expressed genes were identified using a minimum 1.5-fold change and an adjusted p-value ≤ 0.05 after multiple-testing correction. Gene Ontology (GO) enrichment analysis was done using the PlasmodBGO database (Aurrecoechea *et al*., 2009).

### Differential Exon Usage Analysis

For differential exon usage analysis, the reference annotation was prepared using *dexseq_prepare_annotation.py* from the DEXSeq package. Exon-level read counts were subsequently generated from the aligned RNA-seq reads using *dexseq_count.py* (Anders *et al*., 2012). Gene-level read counts were generated using featureCounts from the Subread package (Liao *et al*., 2013) using the BAM files produced by STAR alignment.

### Virulence Gene Family Dynamics Analysis

To assess the dynamics of virulence-associated gene families, the literature was systematically reviewed to identify genes previously implicated in malaria severity (Cheng *et al*., 1998; Lavazec *et al*., 2006; Turner *et al*., 2011). Virulence gene families were extracted from the gene count matrix and analyzed using an in-house R script. Weighted gene co-expression network analysis (WGCNA) was used to assess co-expression and co-regulation patterns among virulence-associated gene families (Langfelder & Horvath, 2008).

### Data Management and Statistical Analysis

Data processing, statistical analysis, and visualization were performed in R version 4.5.2 using Bioconductor packages DESeq2 and DEXSeq. Data wrangling and visualization were done with the tidyverse package.

## Supplementary

See supplementary information

## Data availability

No extra data was generated for this analysis. The code for analysis can be found on our GitHub repository https://github.com/princemensahansah/DEU_DEG_Malaria

## Author contribution

Conceptualization and study design, P.M.A.; Data analysis, P.M.A; literature search and review, P.M.A, W.O.D, A.M; writing—original draft preparation, P.M.A, W.O.D, and A.M.; writing—review and editing, P.M.A, W.O.D, A.M., H.A., and W.A.; formatting and design— P.M.A, W.O.D, A.M., H.A., and W.A.; supervision, W.A. All authors have read and agreed to the published version of the manuscript.

## Acknowledgement

The authors acknowledge Alabama Supercomputer authorities for the computational resources for sequence analysis and mapping

